# Cell wall damage increases macromolecular crowding effects in the *Escherichia coli* cytoplasm

**DOI:** 10.1101/2022.10.03.510584

**Authors:** Theodoros Pittas, Weiyan Zuo, Arnold J. Boersma

**Affiliations:** DWI-Leibniz Institute for Interactive Materials, Forckenbeckstrasse 50, 52074 Aachen, Germany; Institute of Technical and Macromolecular Chemistry, RWTH Aachen University, Worringerweg 1, 52074 Aachen, Germany; Cellular Protein Chemistry, Bijvoet Centre for Biomolecular Research, Faculty of Science, Utrecht University, Utrecht, the Netherlands

## Abstract

The intracellular milieu is crowded with biomacromolecules. Macromolecular crowding changes the interactions, diffusion, and conformations of the biomacromolecules. Changes in intracellular crowding effects have been mostly ascribed to differences in biomacromolecule concentration. However, the spatial organization of these molecules should play a significant role in crowding effects. Here, we find that cell wall damage causes increased macromolecular crowding effects in the *Escherichia coli* cytoplasm. Using a genetically-encoded macromolecular crowding sensor, we see that crowding effects in *E. coli* spheroplasts and Penicillin G-treated cells well surpass crowding effects obtained using hyperosmotic stress. The crowding increase is not due to osmotic pressure, cell shape, crowder synthesis, or volume changes, and therefore not crowder concentration. Instead, a genetically-encoded nucleic acid stain and a small molecule DNA stain show nucleoid expansion and cytoplasmic mixing, which could cause these increased crowding effects. Our data demonstrate that cell stress from antibiotics or cell wall damage alters the biochemical organization in the cytoplasm and induces significant conformational changes in a probe protein.

## Introduction

The bacterial cytoplasm is a spatiotemporal continuum of dynamic interactions of macromolecules, which reflect its physicochemical properties. The total macromolecule concentration has been estimated at 300 mg/mL for *E. coli* ^1,2^. A macromolecule in the cytoplasm that undergoes a conformational change, diffuses, or assembles with other bulky components is affected by the other macromolecules through steric repulsion and nonspecific chemical interactions ^3–6^. Steric repulsion between the macromolecule and other macromolecules decreases the entropy of the system. This entropy can be increased by reducing the volume of the macromolecule(s). For a protein to compress, it needs an internal space where the crowders are excluded so that a (colloidal or osmotic) pressure difference can be built up, i.e., the depletion force. The nonspecific interactions are a sum of steric and chemical components and are weak at the single protein level, but should become more pronounced at the ensemble level in a cell.

In vivo, crowding effects are complicated by the chemical and physical diversity of crowders and their heterogeneous distribution in the cell ^6,7^. Cell components compartmentalize by demixing or preferential interactions in which protein crowders assemble with other biomolecules or membranes. Indeed, diffusion experiments have shown significant spatial heterogeneity in prokaryotes and eukaryotes ^8,9^. Crowders captured in large complexes crowd less effectively because a crowder needs to diffuse to generate the depletion force, and complexation reduces the crowder number density. When crowders do not move, they will cause confinement effects, which are less pronounced until the confinement reaches the shape of a particular macromolecule. Hence, in the compartmentalized cell where biomolecules self-organize at various time and length scales to generate a heterogeneous cytoplasm, the effect of macromolecular crowding may not scale with bulk properties such as biopolymer volume fraction or protein density.

Cells exposed to stresses may see their crowding deviate from optimal levels. Crowding increases are observed when cell volume is reduced by osmotic stress or mechanical pressure, or increased biopolymer synthesis without a volume increase, as reported in some cases for eukaryotes.^10^ In all these cases, it is the biopolymer concentration that changes. Crowding-sensitive probes allow the determination of these rapidly changing crowding effects. These probes are based on diffusion or protein conformation. The former measures viscosity, which, among others, depends on the relative size of the tracer and crowders and is sensitive to large immobile obstacles such as a membrane ^9,11^. Conformationally-sensitive probes rely on compressing a probe protein ^12,13^. A genetically encoded FRET-based probe for macromolecular crowding comprises a pair of two fluorescent proteins that form a FRET pair connected with a linker that facilitates compression in crowded environments, increasing FRET efficiency. The biosensor is sensitive to steric repulsion from crowders and has been applied in buffer, *E. coli*, yeast, and human cell lines in various conditions ^12,13^. Careful choice of the fluorescent proteins allows addressing artifacts such as pH sensitivity or maturation, which is relevant under stress conditions ^14^. This probe showed that crowding effects might deviate from the biopolymer volume fraction; osmotic stress-adapted cells have a lower effective crowding than expected from the biopolymer volume fraction ^15^. In addition, energy depletion reduced crowding effects. Hence, most reported stresses that alter crowding do so through the biopolymer volume fraction, while exceptions exist.

Cell wall damage is particularly harmful to cells, and many antibiotics act on cell walls. Complete or partial cell wall removal with β-lactam antibiotics such as the classical Penicillin G results in cell wall damage-induced dormancy or death in bacteria ^16–21^. Penicillin inhibits murein-specific enzymes (penicillin-binding proteins) and disrupts cross-linking of the peptidoglycan in the cell wall ^16,22,23^. Complete removal of the mechanical support leads to spherical cells named spheroplasts, or protoplasts. Spheroplasts can be generated with lysozyme- or Penicillin G ^24–26^, or result from inhibiting cell shape-governing proteins ^27^. The biopolymer volume fraction in spheroplasts has been determined. One study showed that these cells have the same biopolymer volume fraction as exponentially growing cells ^1^. In contrast, other groups measured an increase in biomacromolecule content (DNA, RNA, and protein) ^25,26,28–32^: Spheroplasts continue to synthesize molecules and can remain viable and regain their usual rod-like shape after spheroplasting ^25,33–35^. Cell wall disruption can lead to a host of downstream effects. On one occasion, it was reported that cell wall damage with Vancomycin reduced the diffusion of a DNA plasmid ^36^, contrasting with various other types of antibiotics. Hence, while cell wall damage is a crucial mechanism for antibiotics, it is unclear whether it affects downstream crowding effects.

Here, using a FRET-based probe for macromolecular crowding, we show that partial or complete cell wall removal strongly increases macromolecular crowding effects. We see that the cytoplasm re-organizes where the nucleoid is less compartmentalized and increased mixing occurs. A better-mixed cytoplasm may increase effective crowding.

## Results

### Spheroplasting increases macromolecular crowding in *E. coli*

We first investigated the effect of spheroplasting in *E. coli* to determine the role of the cell wall on macromolecular crowding effects. We used two variants of our genetically-encoded crowding sensor: the crGE and crGE2.3, which contain mCerulean3/mCitrine and mEGFP/mScarlet-I as FRET pairs, respectively. We quantified the FRET efficiency by dividing the FRET channel over the donor channel. The sensors are distributed homogeneously in the cytoplasm of exponentially growing cells and spheroplasts (Figure 1b). We generated spheroplasts using a lysozyme-based protocol. We found that spheroplasts induce a substantial higher FRET ratio compared to exponentially growing cells (Figure 1c, d) for both the crGE from 0.93±0.05 to 1.11±0.06 (±s.d., n=3) and the crGE2.3 probe from 0.14±0.01 to 0.24±0.04 (±s.d., n=3). pH or maturation artifacts in the FRET ratio from the crGE are opposite to crGE2.3 ^14,53^. Therefore, a FRET increase in both sensors indicates a conformational change to a compressed state.

**Figure 1.**
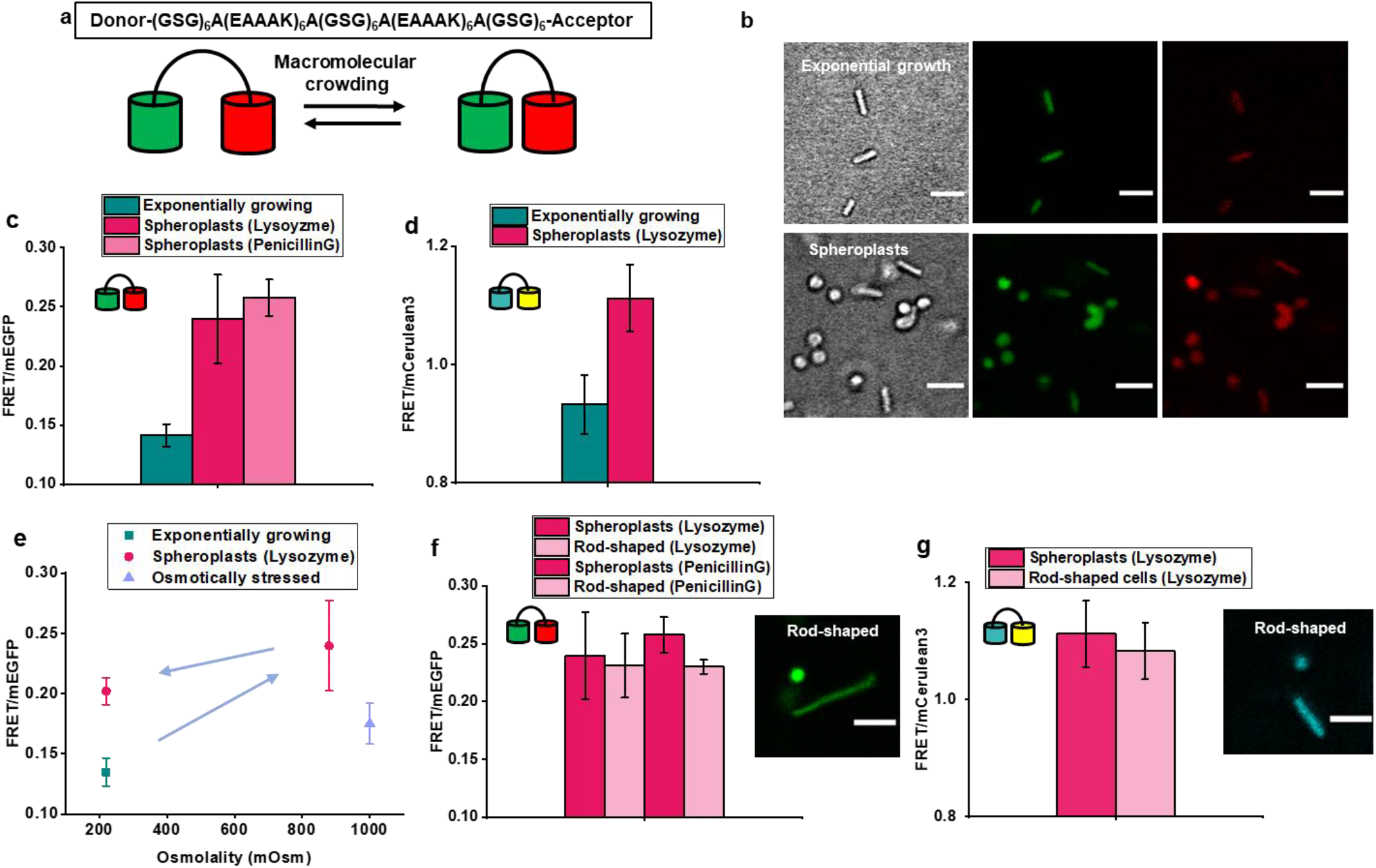
Spheroplast formation increases macromolecular crowding in *E. coli*. (a) The crGE2.3 (mEGFP/mScarlet-I) crowding sensor. Crowding reduces the distance between the acceptor and the donor, increasing FRET efficiency. (b) Representative confocal fluorescence microscopy images of exponentially growing cells and spheroplasts containing the crGE2.3 crowding sensor. The left panels is the brightfield, the middle panels the donor emission, and right panels the acceptor emission upon donor excitation. (c) FRET/Donor emission ratio of exponentially growing cells and spheroplasts created either with lysozyme- or Penicillin G-based protocols. (d) FRET/mCerulean3 for crGE of exponentially growing cells and lysozyme spheroplasts, (e) The effect of osmolality on the crowding in spheroplasts. FRET/mEGFP of exponentially growing cells and osmotically stressed cells with 500 mM NaCl compared to the external osmolality of spheroplasts. Arrows are the order of treatment. (f) comparison of FRET/mEGFP in spheroplasts and rod-shaped cells remaining in the spheroplasting medium. The image depicts typical spherical and rod-shaped cells in the spheroplasting medium. (g) As in (f) but with crGE. Error bars in c-g represent the standard deviation of the average FRET ratios of three independent biological replicates. The size of the scale bar is 4 μm.

To assess the sensitivity to the external osmotic pressure, we returned the spheroplasts stepwise to the growth medium (MOPS minimal medium, ∼220 mOsm) (Figure 1e). We see that the cells show an expected reduction in crowding to a ratio of 0.20±0.01 (±s.d., n=3), albeit this is still higher than cells with an intact cell wall at the same external osmolality. The fluorescence from control cells (without transfected plasmid) was similarly low in exponentially growing cells and spheroplasts, thus excluding any autofluorescence-induced artifacts. We next verified that the crGE2.3 functions in *E. coli* as previously shown in yeast and buffer.^53^ Indeed, we saw an increase in FRET ratio upon a 500 mM NaCl osmotic upshift (∼1 Osm) of exponentially growing cells from 0.14±0.01 to 0.18±0.02 (±s.d., n=3). The ratio after osmotic upshift is lower than upon spheroplast formation at ∼0.8 Osm, indicating the drastic crowding change during spheroplasting (Figure 1e). Finally, we tested whether the increase was related to the lysozyme-based spheroplasting protocol but observed similar FRET ratios when using Penicillin G instead of lysozyme (Figure 1c and Supplementary Figure 1). Hence, spheroplasting increases macromolecular crowding effects.

While the volume decrease of spheroplasts (Figure 2d) would suggest that higher shrinkage induces a higher crowding, we find that incompletely spheroplasted cells that retain the rod shape give the same ratios as the fully spheroplasted cells (Figure 1f, g). This surprising result shows that cell volume, shape, and the mechanical constraints of the cell wall may not play a role in the spheroplast crowding increase.

**Figure 2.**
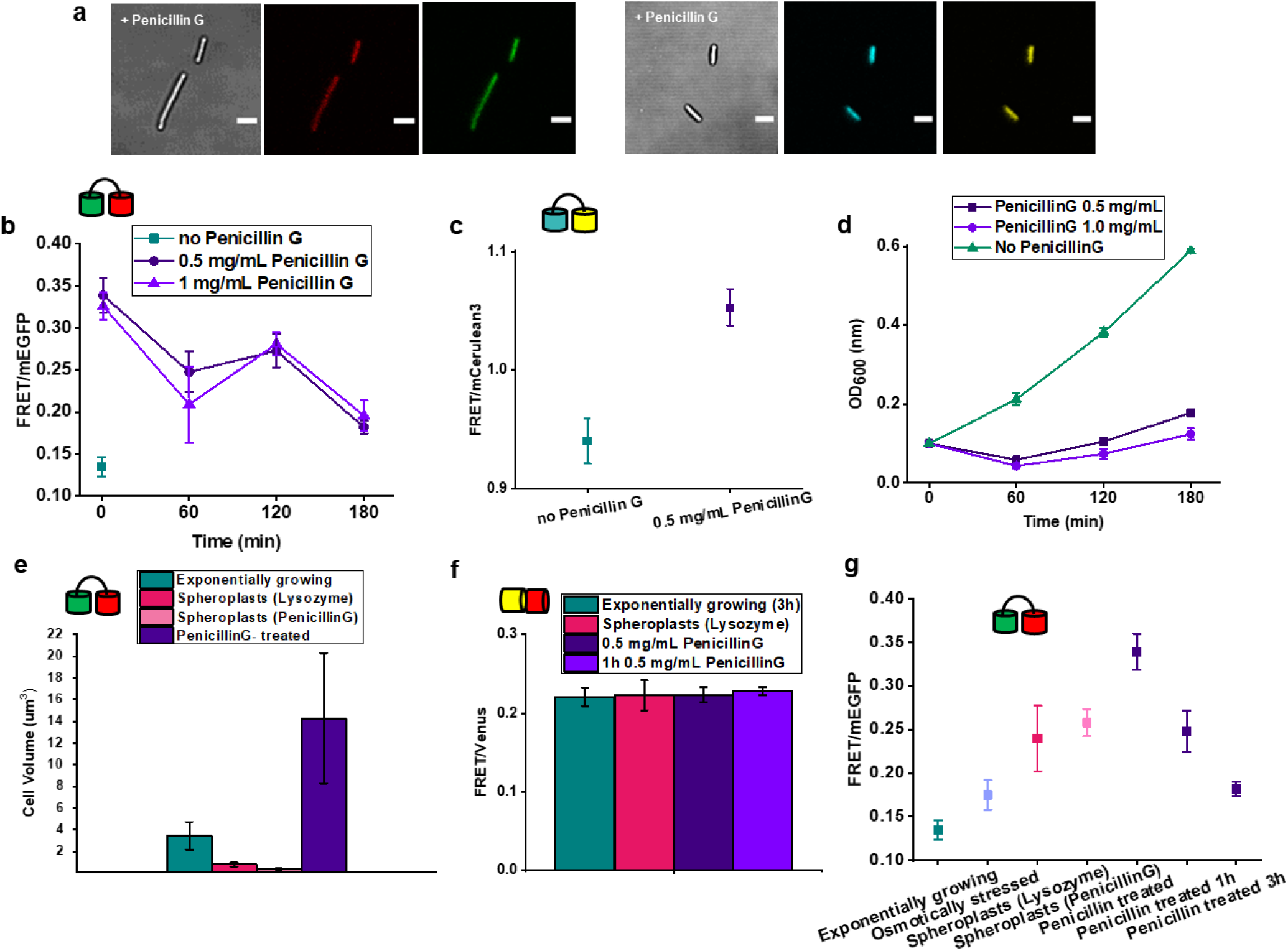
Penicillin G increases macromolecular crowding. (a) fluorescence confocal microscopy images after Penicillin treatment of *E. coli* BL21 with crGE3.2 (left three panels) and crGE (right three panels). Displayed are brightfield, donor, and acceptor channels after donor excitation. (b) Time dependence of FRET/mEGFP treated with 0.5 or 1.0 mg/mL Penicillin. (c) FRET/mCerulean3 ratios for cells treated with 0.5 mg/mL Penicillin G for 1 minute. (d) Culture growth as optical density (OD_600_) dependence on Penicillin G. (e) Cell volume of exponentially growing cells, spheroplasts and PenicillinG-treated cells. (f) FRET/mVenus ratios of the mVenus/mCherry construct in exponentially growing cells, spheroplasts, and Penicillin G-treated cells, (g) Comparison of FRET ratio changes in various stress conditions. Error bars represent the standard deviation of the average of three independent biological replicates. The size of the scale bar is 4 μm.

### Cell wall damage causes a macromolecular crowding increase

To verify whether cell wall damage without spheroplasting is sufficient to increase the macromolecular crowding as we see for incompletely spheroplasted cells, we treated exponentially growing cells with Penicillin G and monitored crowding in time. We chose the two subinhibitory concentrations of Penicillin G used for spheroplasting ^25^ to treat our cells. Penicillin G temporarily halted growth, and cell growth resumed two hours after the treatment (Figure 2c). If stopping cell growth would result in a build-up of synthesized macromolecular crowders, the crowding should gradually increase. Unexpectedly, the FRET ratio increased within a minute to 0.34±0.02 (±s.d., n=3) after adding 0.5 mg/mL Penicillin G (Figure 2a), which is higher than the spheroplasts. We tested the crGE probe, which also showed an immediate increase in FRET ratio to 1.05±0.02 (±s.d., n=3) (Figure 2b), which excludes pH effects. We see that the ratios remain high and decrease slowly over three hours of adaptation in MOPS minimal medium containing Penicillin G to approach the FRET ratios of the spheroplasts. During the decrease in crowding effects, the OD increases slowly (Figure 2d). To completely exclude optical or fluorescent protein artifacts in these stress conditions, we tested a mVenus-mCherry construct without a linker ^37^ to prevent intramolecular FRET changes. We found that the FRET ratios of spheroplasted and Penicillin-treated cells are the same as exponentially growing cells (Figure 2e). Hence, the FRET increase is due to a compressed conformation of the crowding sensor, instantaneously induced by the action of Penicillin G on the cell wall.

The 1-min timeframe is too short for biopolymer synthesis to significantly increase crowding, considering that the cell maintains a 60-minute doubling time. Therefore, we assessed whether the crowding increase is due to a decrease in cell volume. We determined the cell volume before perturbation from microscopy images of the crGE2.3 probe in the cytoplasm. We find a cell volume of 4.1±0.2 μm^3^, which decreases to 0.8±0.2 μm^3^ and 0.38±0.01 μm^3^ for lysozyme and Penicillin G-based spheroplast formation, respectively (Figure 2d). In contrast, cells that remain rod-shaped have the same appearance as before spheroplasting. The volume of Penicillin G-treated cells increased steeply to 14±6 μm^3^ one minute after treatment due to the cell wall damage. Therefore, the cytoplasmic volume does not correspond to the crowding increase. The increase in FRET due to Penicillin G also offers an explanation for the increase in crowding in spheroplasts, as cell wall damage appears to be sufficient to increase crowding.

### Cytoplasmic mixing is a potential cause for the crowding increase

We next aimed to determine a potential cause for the crowding increase. Crowding effects depend not only on the absolute concentration of biomacromolecules or the biopolymer volume fraction in the entire cell but also on the macromolecular organization. For example, altering crowder distribution or their diffusion rate by complexation in weakly assembled large immobile complexes would generate less crowded areas where a sensor resides, and the effective crowding decreases. Moreover, a well-mixed cytoplasm would induce higher crowding as there are more collisions with crowders. The nucleoid forms a compartment in *E. coli* and is one of the main organizers of the cell: a compact nucleoid will take up less space, reducing the effect of protein-based crowders. Indeed, the nucleoid/cytoplasm ratio has been shown to affect cell biophysical properties strongly ^38^. We therefore assessed the properties of the nucleoid to test the hypothesis that the macromolecular organization changes significantly upon cell wall damage, which could increase crowding.

We first used a genetically-encoded polynucleotide stain based on (KWK) repeat units fused to GFP to image the nucleoid.^55^ We used a derivative based on two (KWK)_2_ sequences for divalent binding. This probe has been shown to bind the nucleoid of *E. coli*, although these peptides also bind RNA in vitro ^39,40^. We imaged the KWK probe localization by confocal fluorescence microscopy after 4-5h expression with 100 μM IPTG. The probe is excluded from the cell poles in MOPS minimal medium and follows the location of the nucleoid (Supplementary Figure 2). When we use LB medium, we see fluorescent foci in ∼70% of the cells (Figure 3a, Supplementary Figure 3, 5), with the majority localizing at the poles. The poles are usually associated with the ribosomes and not the nucleoid, possibly becoming a binding partner at high expression levels in LB, which does not occur at lower KWK expression levels in MOPS. Upon spheroplasting, we see complete fluorescence mixing in the cytoplasm of MOPS and LB-grown cells, indicating a reduced organization and increased mixing (Supplementary Figures 3-5). For the addition of Penicillin G to exponentially growing cells, we see rapid dispersion of the KWK foci in about 75% of the cells (Figure 3a, d, Supplementary Figure 3-5). After Penicillin G, the probe occupies the entire cell, including the pole regions. Non-spheroplasted cells in spheroplasting medium showed a similar spread pattern to the Penicillin-treated ones, thus indicating possible similarities between these two cellular states. Cells grown in MOPS medium did not have the foci that Penicillin G could disperse, and the probe remained dispersed (Supplementary Figure 2). Hence, the genetically-encoded polynucleotide-binding probe spreads throughout the cytoplasm upon cell wall damage.

**Figure 3.**
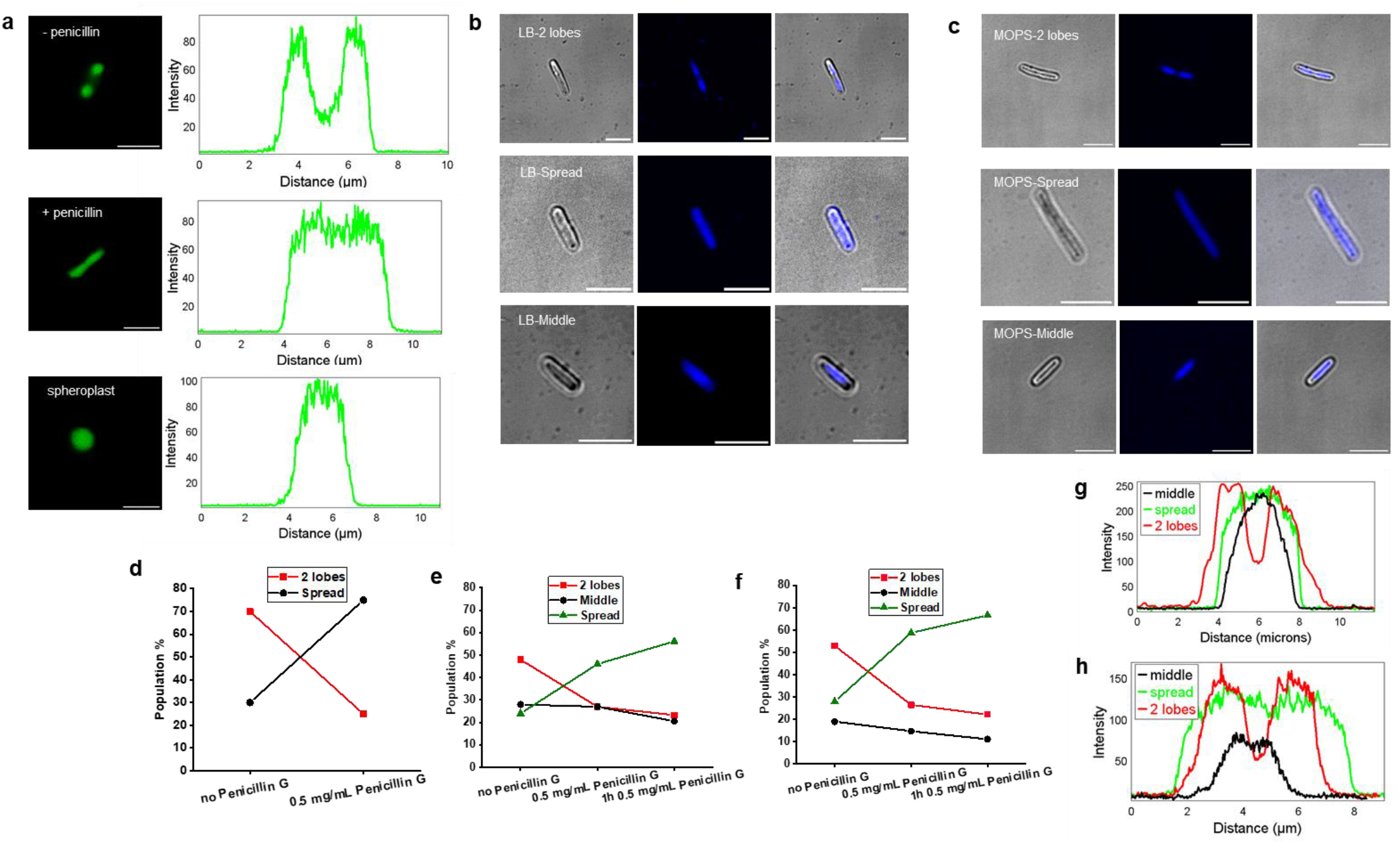
The nucleoid is more spread upon spheroplasting or Penicillin treatment. (a) Fluorescence confocal microscopy images of *E. coli* BL21 expressing the KWK probe showing a difference in the distribution in exponentially growing cells in LB medium, Penicillin G-treated cells (1 minute after adding 0.5 mg/mL Penicillin G), and spheroplasts. Right panels are the corresponding fluorescence intensity over the longest axis in the bacteria. (b) Fluorescence confocal microscopy images of DAPI-stained nucleoids in LB medium show three distinct categories. (c) as in (b), cells were grown in MOPS medium. (d) Increase the percentage of cells with a spread nucleoid conformation upon Penicillin G addition. (e) as in (d) where DAPI stain is used in LB medium, showing a similar pattern. (f) as (e) with MOPS medium. (g) DAPI fluorescence intensity along the longest axis of the bacteria for LB medium. (h) as in (g) with MOPS medium. >35 cells per condition were analyzed.

To better map the consequences of cell wall damage, we stained the nucleoid selectively with DAPI. Exponentially growing cells can be grouped into three populations; those with two lobes, those with one mid-lobe, and those with a spread nucleoid (Figure 3c, e, Supplementary Figure 5). These likely correspond to the cell cycle stage: localization of DNA in 2 lobes is the characteristic pattern of the end of the replication cycle and beginning of division. In MOPS and LB, the 2-lobed configuration is twice as prevalent as the other configurations. The similarity between MOPS and LB shows that the KWK foci in LB are not due to nucleoid staining but are more likely rRNA or mRNA staining. Within 1 minute after the addition of Penicillin G, we see that the spread nucleoid state is twice as prevalent as the lobed states. The nucleoid spreads throughout the entire cell, including the poles. The spread state is even more dominant after one-hour incubation with Penicillin G. These findings correspond to the dispersion of KWK foci and KWK spreading through the cytoplasm upon the addition of Penicillin G. An increased mixing was also observed in spheroplasts, where we used DRAQ5 dye to stain the nucleoid (Supplementary Figure 6). Hence, cell wall damage homogenizes cytoplasmic content, as demonstrated by the nucleoid expansion and dispersion of an oligonucleotide-binding probe. These mixing effects offer a potential route to higher macromolecular crowding effects (Figure 4).

**Figure 4.**
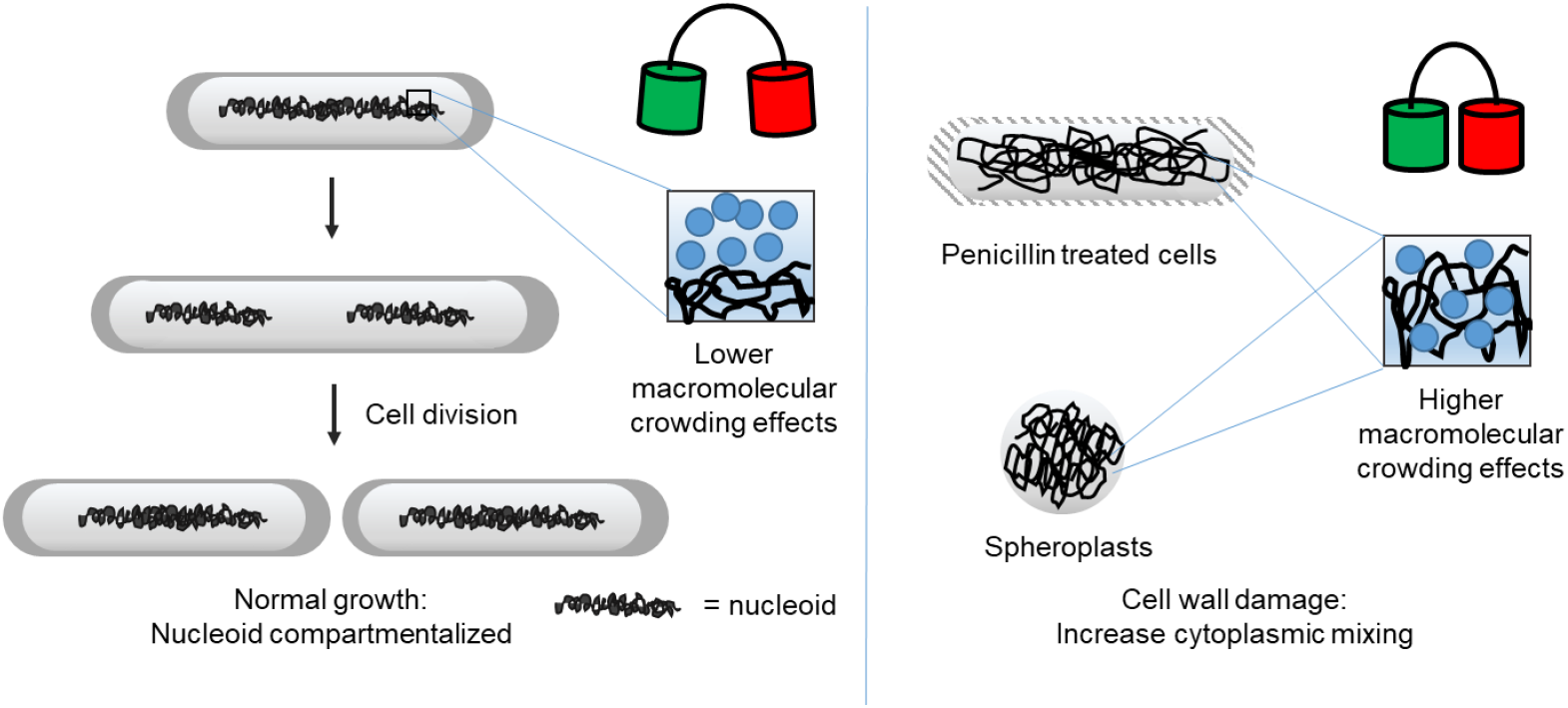
Schematic representation of nucleoid expansion with cell wall damage. Left panel: exponentially growing cells that divide and maintain a more compartmentalized cytoplasm. Right panel: partial (Penicillin G) or complete (spheroplasts) cell wall damage inhibits division and provides cells with a more expanded nucleoid. The more homogenized interior could lead to higher macromolecular crowding effects.

## Discussion

This study used a genetically-encoded FRET biosensor to quantify macromolecular crowding effects in *E. coli* cells with a partly- or fully-removed cell wall. We find that a) crowding effects in spheroplasts and Penicillin-treated cells surpass the ones measured in osmotically stressed cells, b) the observed effects are irrespective of cell shape, external osmolality, biopolymer synthesis, and cell volume, and c) cell wall damage possibly results in cytoplasmic homogenization and changes in nucleoid organization.

Our findings rely on sensors to measure macromolecular crowding. These sensors have been validated in different cell types, including during osmotic upshift in *E. coli* ^12,13^. The sensors measure steric interactions, thus excluding interactions with proteins of the bacterial cytoplasm, while most other proteins would respond to additional nonsteric interactions under crowded conditions^47,48^. The sensors give information on the crowding effect at the 5-10 nm scale, which is in the FRET range. The sensors thus measure crowders or confinements in a similar size range, which is a typical size for an intracellular protein. The readout is not due to fluorescent protein effects. Therefore, beyond reasonable doubt, the sensor is highly compressed upon cell wall damage, showing drastic physiological changes. This compression is indicative of an effective macromolecular crowding increase at the size range of a typical protein.

Up to date, crowding effects have most frequently been correlated with volume changes and protein concentration ^1,10,13^. Also the crowding sensor reported crowding values corresponding to cell volume changes upon osmotic stress. However, in previous work, we noticed that cells adapted to osmotic stress or energy-depleted cells displayed reduced effective crowding, showing that crowding effects may not scale with the biopolymer volume fraction in the entire cytoplasm ^15^. Here we show the crowding can also greatly increase independent from the cell volume. In spheroplasts, the longer preparation time could in principle also increase crowding due to continued biopolymer synthesis that was measured before ^25,31^, and these cells are much smaller. However, the Penicillin-treated cells and spheroplasts are linked by cell wall damage: Even cells that are incompletely spheroplasted give the same results as Penicillin-treated cells. The Penicillin-G treated cells have a higher volume immediately after treatment, and therefore the concentration of macromolecular cannot increase but their crowding immediately after treatment is higher than spheroplasts. Therefore, additional biopolymer synthesis and cell volume effects may play a role, but this is not the main driver of the high crowding upon cell wall damage.

The link between the cell wall and crowding that we see is not straightforward. The cell wall regulates the turgor pressure and maintains surface to mass ratio ^41^. However, the short timeframe excludes a discrepancy between halted cell wall growth and cell volume and mass. On the other hand, many of the proteins involved in cell division are associated with cell wall maintenance and coordinate the segregation of DNA during cell division ^26,42,43^. DNA is a prominent organizer in the cell, and its perturbation would have consequences on the physical-chemical properties of the cell. For example, under conditions that favor growth, the cytoplasm is a bad solvent for the nucleoid ^38^, giving it its specific architecture and organization. In these conditions, DNA is withheld away from cell΄s end caps by entropic forces, while ribosomes and RNA species are more common in the end caps ^44^. In healthy cells, DNA is dynamic and replication begins in the mid-cell and in the intermediate stage of the cycle DNA molecules spread towards the poles. At the final stage of replication, the chromosome segregates in two lobes, and the daughter cell is detached ^42,45,46^. Perturbation of these processes through the cell division machinery, improving the solvent quality of the cytoplasm, or altered transcription/translation may alter the DNA shape, conformation, and its mixing with the cytoplasm. We see in both spheroplasting and Penicillin treatment that the nucleoid is expanded. These cell wall perturbations are highly suitable for our sensors, as they do not stop transcription and translation, which would alter the fluorescence properties of our sensors by generating maturation artifacts.

How can the expansion of the nucleoid lead to crowding changes? Of note is that the crowding increase is exceptionally high, much higher than the osmotic upshift, suggesting a dramatic change in cytoplasmic material properties. Indeed, it has been shown that increased nucleoid size reduces the diffusion of larger particles.^38^ While several explanations can be put forward in lieu of a clear theory or relevant in vitro experiment, we deem it most likely that the number of free (or less crowded) spaces is reduced by a better distribution of all the cell components. This would lead to more collisions between the crowders and the sensor, and a higher effective crowding readout. These collisions may be with diffusing crowders or with the walls in confined spaces. A better distribution could be caused by mixing in dense protein regions, such as the RNA and protein density at the cell poles or release of nucleoid associated proteins. Also the sensor may be better distributed, providing a better average crowding without being crowded out of denser regions.

A change in the macromolecular crowding and physicochemical properties of the bacterial cytoplasm may be a more common response to antibiotics: it was observed that Vancomycin, also a cell wall inhibitor, decreases the motility of DNA and cytosol ^36^. Moreover, nucleoid spreading is seen for other antibiotics, such as rifampicin and cecropin A, where nucleoid expands and mixes with ribosomes ^49–51^. In addition, LL37 also makes the nucleoid appear more diffuse, albeit this likely functions through cross-linking DNA ^52^. While these mechanisms may not be the same as described here and may not have the same effect at the protein scale, they induce large cytoplasmic changes and would alter crowding properties. This re-organization of biomolecules would have a plethora of downstream effects affecting biomolecule biogenesis, altering metabolic kinetics, and reducing cell growth, exacerbating the effect of the antibiotic.

## Materials and methods

### Growth conditions of *E. coli* cells

We used *E. coli* BL21(DE3) for every experiment. The synthetic genes crGE2.3 ^53^ and VC^37^ in the plasmid pRSET-A were codon-optimized for *E. coli* and obtained from GeneArt. Bacteria were transformed with crGE2.3 (meGFP-mScarlet-I) or the crGE (mCerulean3-mCitrine) crowding sensor.^13^. A single colony of freshly transformed cells was picked and grown in overnight incubation in (5-6 mL) MOPS minimal medium^54^ with 50 μg/mL ampicillin (TCI) at 37 ^o^C and 180 *rpm* agitation. The cell cultures were diluted to an OD_600_ of 0.03 in fresh MOPS (10 mL) at 100 mL Erlenmayer flasks and grown at 30 ^o^C and 180 *rpm* agitation until they reached an OD_600_ of 0.1-0.3. The sensor was expressed sufficiently without an inducer. Fluorescent cells were mixed with *E. coli* BL21 cells without plasmid of the same growth phase just before imaging. Experiments with spheroplasts, osmotic stress, and Penicillin were conducted subsequently.

### Osmotic stress experiments

For osmotic stress experiments we used 500 mM NaCl, as described also elsewhere ^13^. Briefly, 1 mL of *E. coli* in the exponential growth phase was pelleted at 5000 x *rpm* for 5 min and re-suspended in MOPS medium supplemented with 500 mM NaCl, without any potassium sources or glucose, to avoid adaptation of the cells. The samples were immediately mounted on glass microscopy slides and imaged.

### Preparation of *E. coli* spheroplasts with lysozyme

We created *E. coli* BL21 (DE3) spheroplasts following literature, with minor modifications ^24^. 1 mL of the cell suspension in the exponential growth phase was pelleted at 3000 x *g* for 1 min. The pellet was re-suspended in 500 μL 0.8 M sucrose solution. Then we added the following solutions to the cell aliquots: 30 μL Tris-HCl (pH 8.0), 24 μL 0.5 mg/mL lysozyme, (∼20 μg/mL final concentration), 6 μL 5 mg/mL DNase (∼50 μg/mL final concentration), and 6 μL 125 mM EDTA-NaOH (pH 8.0) (∼1.3 mM final concentration). Incubation of the sample for 10 min at 25 °C followed and 100 μL of STOP solution (10 mM Tris·HCl, pH 8, 0.7 M sucrose, 20 mM MgCl_2_) were added to terminate the digestion. The cell suspension was pelleted at 5000 x *rpm* for 5 min and re-suspended in 50-70 μL of the spheroplasting mixture. 15 μL of the sample were mounted on glass microscopy slides and the creation of *E. coli* spheroplasts was evaluated, using confocal laser scanning microscopy. For the adaptation of spheroplasts to a lower osmolality medium, liquid cultures of spheroplasts were diluted to an osmolality equal to our MOPS medium (220 mOsmol/kg), by addition of milliQ water.

### Preparation of *E. coli* Spheroplasts with Penicillin G

We created *E*.*coli* BL21 (DE3) spheroplasts as described elsewhere, with minor modifications.^25^ Cell suspension in the exponential growth phase was pelleted at 5000 x *rpm* for 5 min and resuspended in 10% w/v sucrose solution. Then we added the following solutions: 0.5 mg/mL of sodium salt of PenicillinG and 0.2% w/v MgSO_4_. After ∼1h incubation, 15 μl of cell suspension were mounted on a glass microscopy slide to evaluate the creation of *E. coli* spheroplasts.

### Penicillin G-treated *E. coli*

After the liquid cell cultures had reached the exponential growth phase they were incubated in MOPS supplemented with 0.5 mg/mL or 1 mg/mL Penicillin G. We selected antibiotic concentrations that were previously used for creating bacterial spheroplasts ^25,33^. Samples were collected immediately, 1h, 2h, and 3h after antibiotic treatment and pelleted at 5000 x *rpm* for 5 min and then 15 μL sample was mounted on glass microscopy slides for imaging.

### KWK probe

The KWK probe was designed based on literature.^55^ The gene was synthesized by W. Zuo and was cloned into the plasmid pRSET-A. *E. coli* BL21(DE3) containing the corresponding plasmid was grown in LB medium following the usual protocol. Protein synthesis was induced with 100 μM IPTG at an OD_600_ of 0.1 at 30 °C for 5-6 h. The cells were mounted on a glass slide and were imaged.

### Nucleoid staining

At an OD_600_ of 0.1 we added 10 μg/mL DAPI^56^ or 10 μM DRAQ5 (final concentration) and allowed the cells to grow for ∼ 2h. Then we created spheroplasts or treated cells with penicillin and used microscopy to evaluate nucleoid localization.

### Microscopy settings & analysis

For imaging *E. coli* cells, glass microscopy slides were mounted on a Leica TCS SP8 laser-scanning confocal microscope. The crGE2.3 sensor was excited using a 488-nm Argon laser and the emission was split into a 500-550 nm and a 580-700 nm intensity channel. The mCerulean3-mCitrine crowding sensor was excited using a 405-nm LED laser, and the emission was split into a 465-505 nm channel and a 525-600 nm intensity channel. The KWK sensor was excited using a 488 nm Argon laser and the emission was set into a 510-550 nm channel. DAPI dye was excited using a 405-nm LED laser and the emission was split into a 430-550 nm channel. DRAQ5 dye was excited using a 510 nm Argon laser and the emission was set into a 600-700 nm channel.

Image analysis was performed with the use of ImageJ ^57^, an open source scientific image processing program. We plotted the FRET channel intensity versus the Donor channel intensity for every FRET-based sensor and under all experimental conditions described in this work. The data were fitted in a linear equation using the least-squares approach and the slope as the average FRET ratio as described previously ^12^.

Fluorescence intensity versus distance plots were made for straight, non-dividing cells using the plot profile option in ImageJ. Cell heat maps were made with Origin 2018b.

## Aknowledgements

The work was funded by the ERC Consolidator Grant (PArtCell; no. 864528) the Netherlands Organization for Scientific Research (NWO Vidi; 723.015.002).

## Contributions

P.T. and A.J.B. conceived the project. A.J.B. and P.T. designed the experiments and T.P. performed the experiments and analyzed the data. W.Z. designed and cloned the KWK sensor. T.P. and A.J.B. co-wrote the paper.

## Supporting Information

**Supplementary Figure 1.**
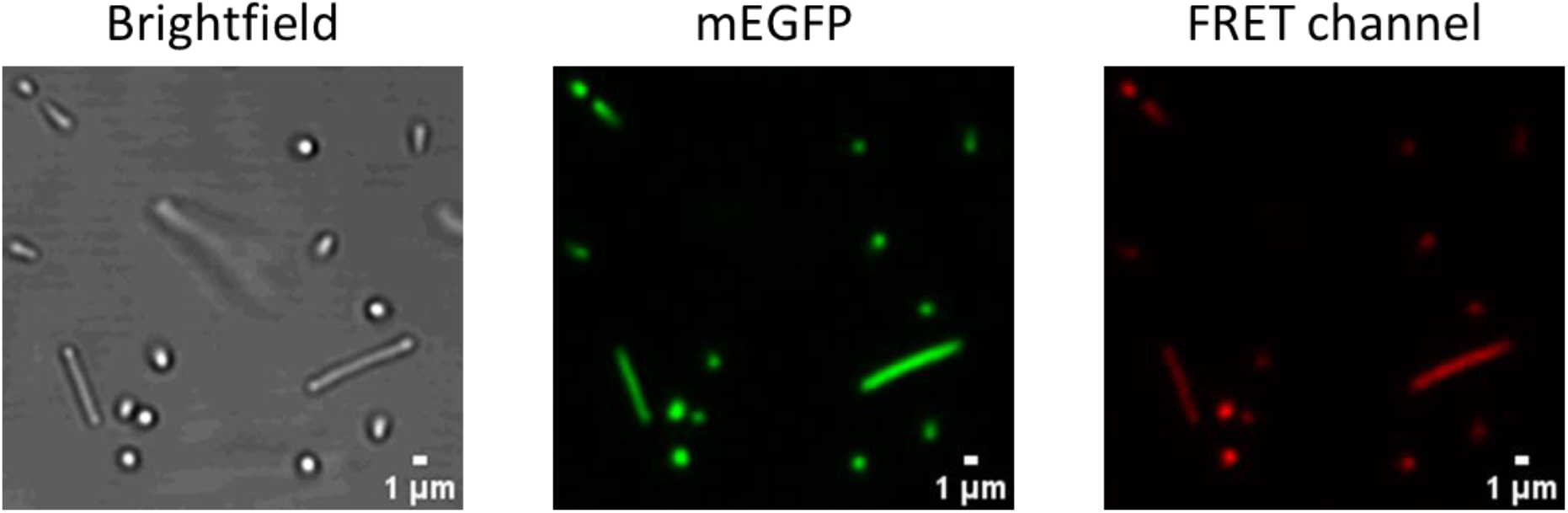
Representative images of spheroplasts created with Penicillin G and transformed with the crGE2.3 (mEGFP-mScarlet-I) crowding sensor.

**Supplementary Figure 2.**
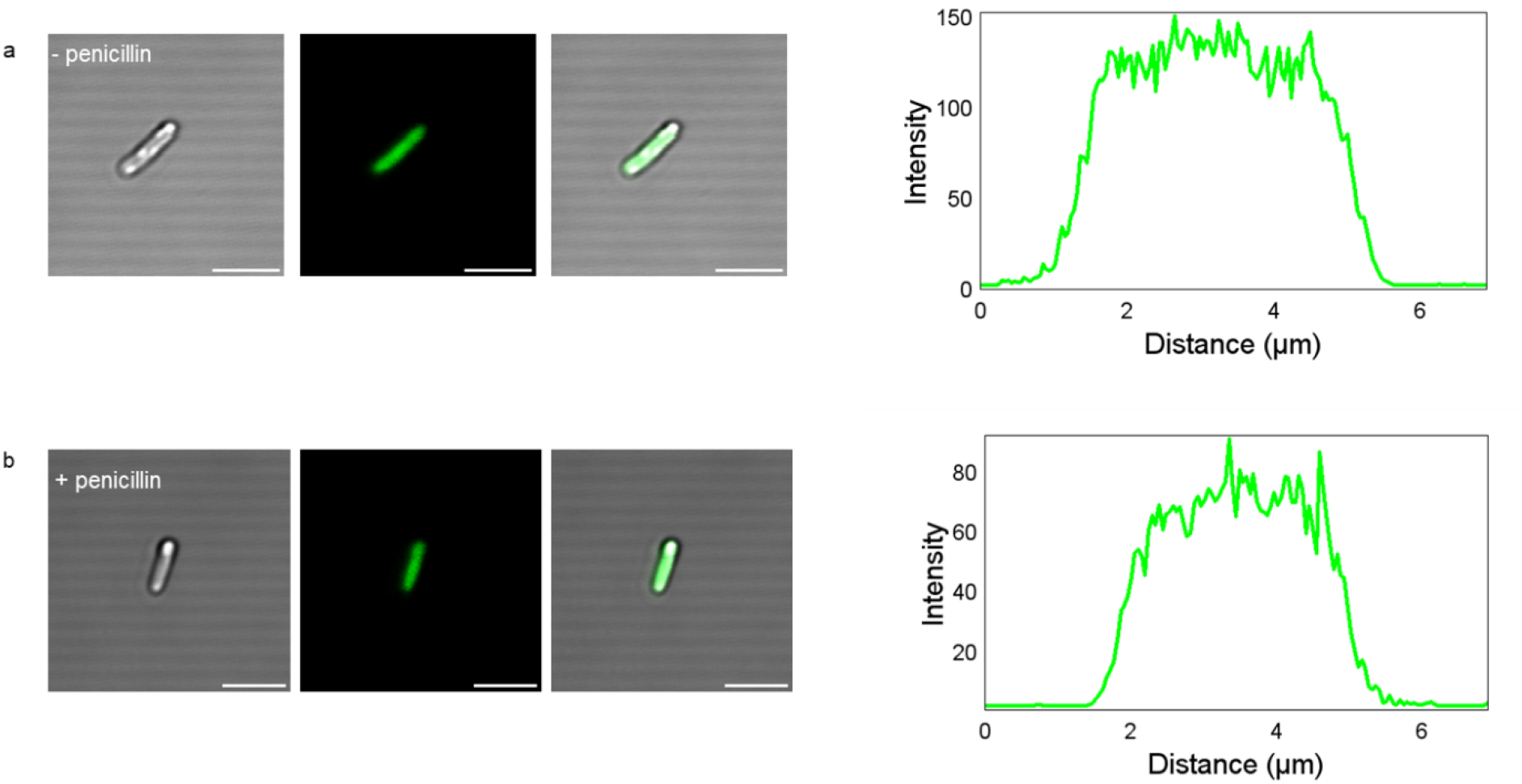
Representative images and localization patterns of the KWK sensor in exponentially growing cells in MOPS growth medium, without penicillin and 1 min after treatment with 0.5 mg/mL Penicillin G.

**Supplementary Figure 3.**
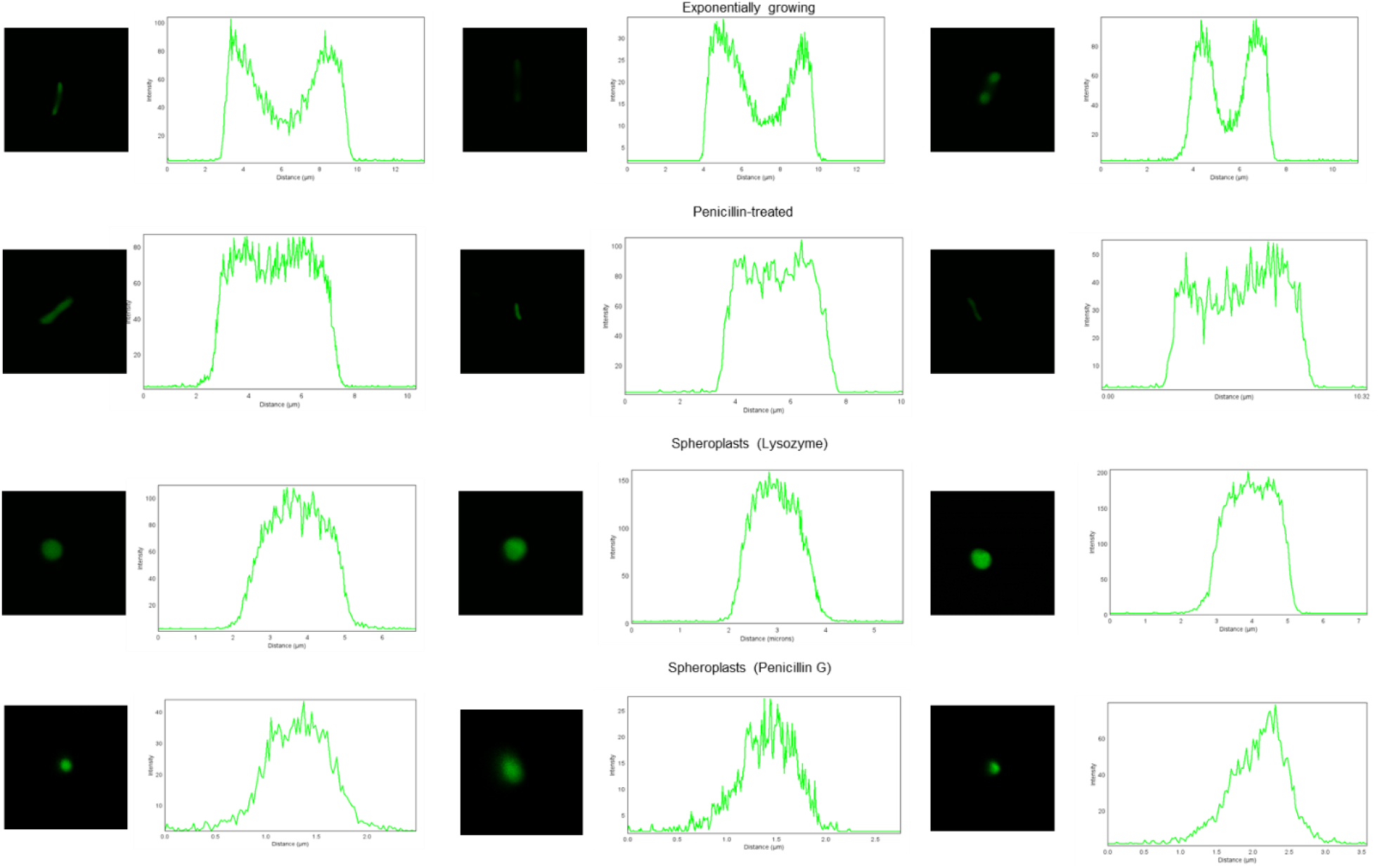
Representative images and localization patterns of the KWK sensor in exponentially growing cells in LB growth medium, spheroplasts, and cells 1 min after treatment with 0.5 mg/mL Penicillin G.

**Supplementary Figure 4.**
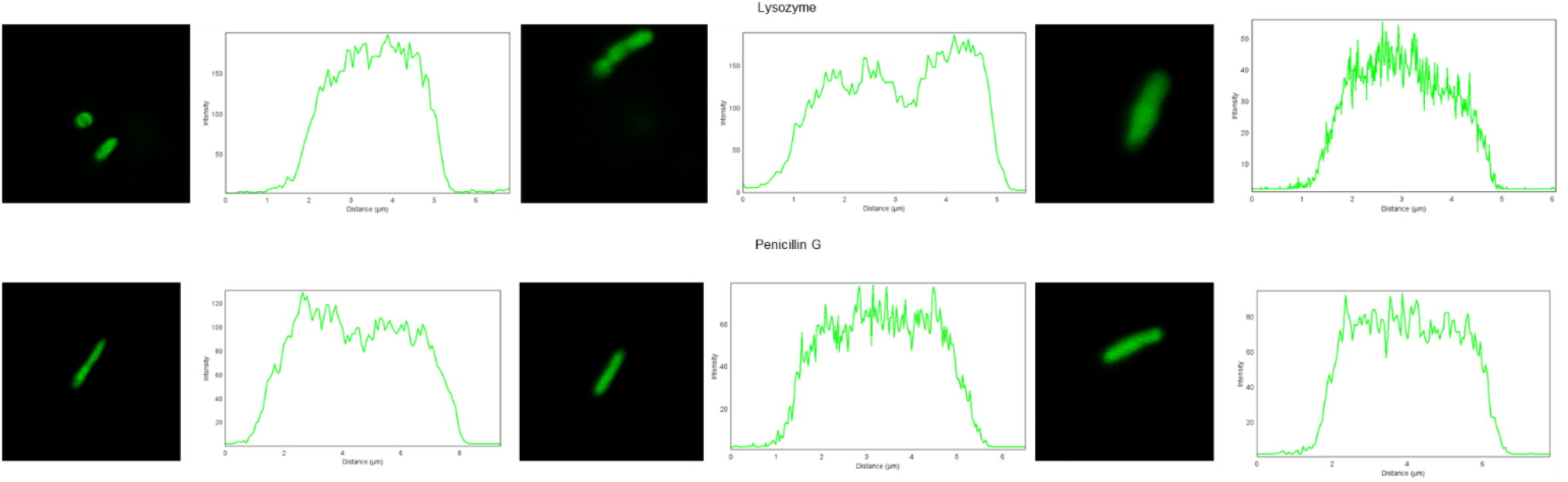
Representative images and localization patterns of the KWK sensor in rod-shaped cells in the spheroplasting medium.

**Supplementary Figure 5.**
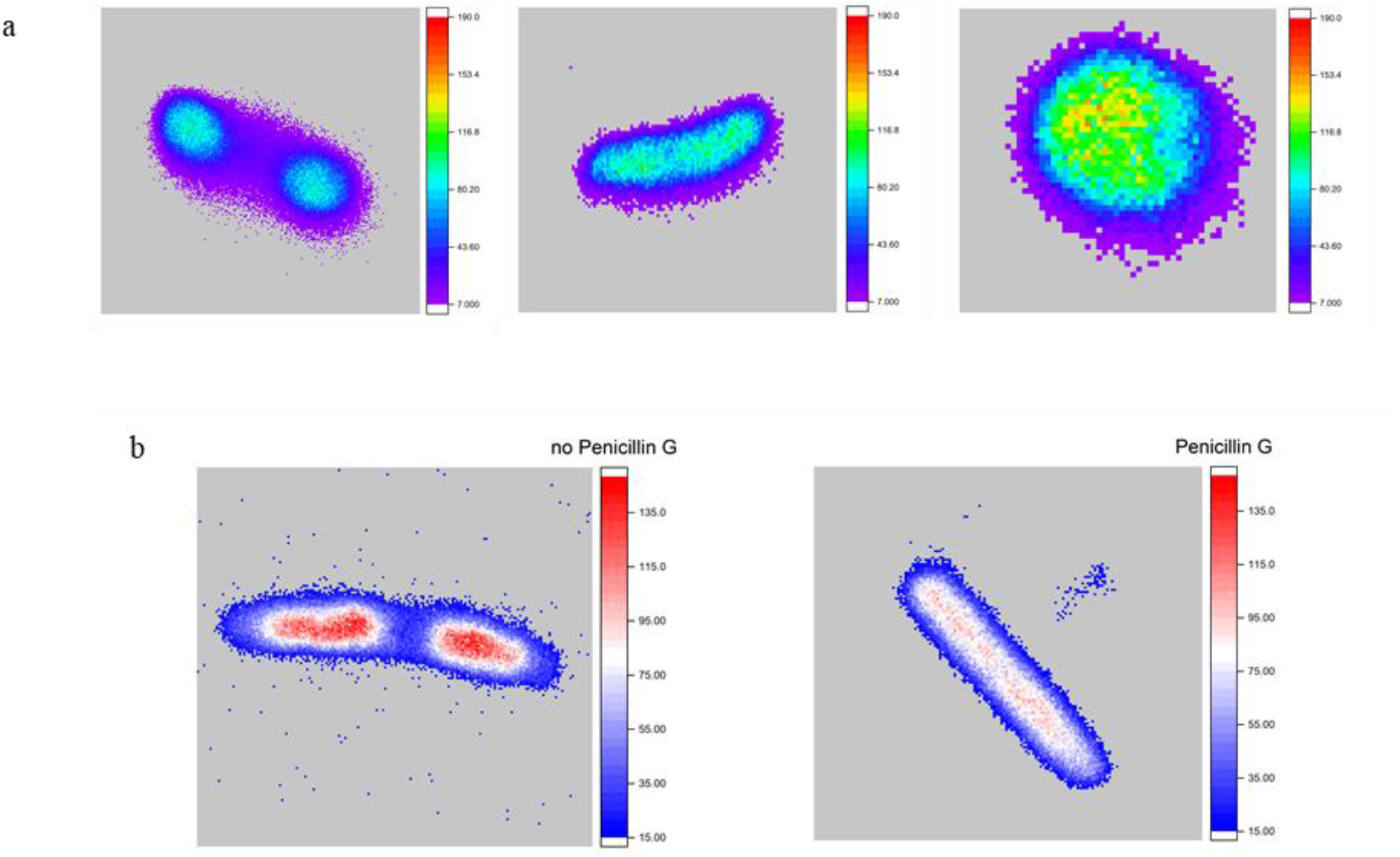
Heat maps depicting the localization of (a) KWK in exponentially growing cells, Penicillin G-treated cells, and spheroplasts, (b) nucleoid-staining dye, DAPI, in exponentially growing cells and cells after 1min of treatment with 0.5 mg/mL of Penicillin G.

**Supplementary Figure 6.**
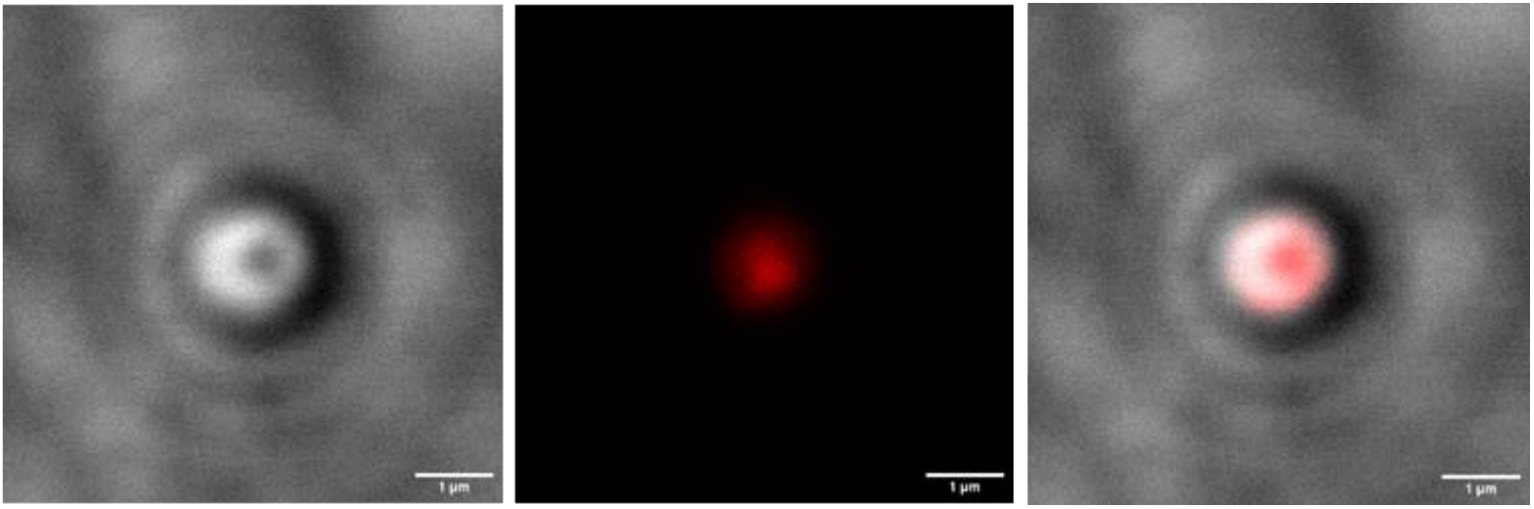
Representative image of nucleoid-staining dye, DRAQ5 in spheroplasts (Lysozyme).

### DNA Sequences

#### CrGE2.3 (mEGFP/mScarlet-I)

ATGAAAGGTGAAGAACTGTTTACCGGTGTTGTTCCGATTCTGGTTGAACTGGATGGTGACGTTAATGGTCACAAATTTTCAGTTAGCGGTGAAGGCGAAGGTGATGCAACCTATGGTAAACTGACCCTGAAATTTATCTGTACCACCGGCAAACTGCCGGTTCCGTGGCCGACACTGGTTACCACACTGACCTATGGTGTTCAGTGTTTTAGCCGTTATCCTGATCACATGAAACAGCACGATTTTTTCAAAAGCGCAATGCCGGAAGGTTATGTTCAAGAACGTACCATCTTCTTCAAAGATGACGGCAACTATAAAACCCGTGCCGAAGTTAAATTTGAAGGTGATACCCTGGTGAATCGCATTGAACTGAAAGGCATCGATTTTAAAGAGGATGGTAATATCCTGGGCCACAAACTGGAATATAATTATAATAGCCACAACGTGTACATCATGGCCGACAAACAGAAAAATGGCATCAAAGTGAACTTCAAGATCCGCCATAATATTGAAGATGGTTCAGTTCAGCTGGCCGATCATTATCAGCAGAATACCCCGATTGGTGATGGTCCGGTTCTGCTGCCGGATAATCATTATCTGAGCACCCAGAGCAAACTGAGCAAAGATCCGAATGAAAAACGCGATCACATGGTGCTGCTGGAATTTGTTACCGCAGCAGGTATTACCTTAGGTATGGATGAACTGTATAAAGGATCCGGTGGTAGCGGTGGTTCAGGTGGTAGTGGCGGTAGTGGTGGCAGCGGTGCAGAAGCAGCAGCAAAAGAAGCCGCTGCCAAAGAAGCGGCAGCGAAAGAGGCTGCCGCAAAAGAGGCAGCAGCGAAAGAAGCAGCGGCTAAAGCAGGTTCAGGCGGAAGCGGAGGCAGTGGTGGATCAGGCGGATCTGGTGGCTCAGGTGCCGAGGCAGCAGCAAAAGAGGCAGCTGCTAAAGAGGCTGCTGCAAAAGAAGCAGCCGCAAAAGAGGCAGCGGCAAAAGAAGCCGCAGCAAAAGCAGGTAGTGGTGGAAGTGGCGGTTCCGGTGGCTCTGGTGGAAGCGGTGGCTCCGGAgagctcGTTAGTAAAGGCGAAGCAGTTATTAAAGAATTTATGCGCTTCAAAGTGCACATGGAAGGTAGCATGAATGGCCATGAATTTGAAATCGAAGGTGAAGGTGAGGGTCGTCCGTATGAAGGCACCCAGACCGCAAAACTGAAAGTTACCAAAGGTGGTCCGCTGCCGTTTAGCTGGGATATTCTGAGTCCGCAGTTTATGTATGGTAGCCGTGCATTTATCAAACATCCGGCAGATATCCCGGATTATTACAAACAGAGCTTTCCCGAAGGTTTTAAATGGGAACGTGTGATGAATTTTGAGGATGGTGGTGCAGTTACCGTTACACAGGATACCAGCCTGGAAGATGGCACCCTGATCTATAAAGTTAAACTGCGTGGCACCAATTTTCCGCCAGATGGTCCTGTTATGCAGAAAAAAACCATGGGTTGGGAAGCAAGCACCGAACGTCTGTATCCTGAAGATGGCGTTCTGAAAGGTGATATCAAAATGGCACTGCGTCTGAAAGATGGTGGTCGTTATCTGGCAGATTTCAAAACCACCTACAAAGCCAAAAAACCGGTTCAGATGCCTGGTGCATATAATGTTGATCGCAAACTGGATATCACCAGCCATAATGAAGATTATACCGTGGTGGAACAGTATGAACGTAGCGAAGGTCGTCATAGTACCGGTGGCATGGATGAATTATACAAAGGTGGCACCTAA

#### crGE (mCerulean3/mCitrine)

ATGCATCATCATCACCATCATGTGAGCAAAGGTGAAGAGCTCTTTACCGGTGTTGTTCCGAT

TCTGGTTGAACTGGATGGTGACGTTAATGGTCACAAATTTTCAGTTAGCGGTGAAGGCGAA

GGTGATGCAACCTATGGTAAACTGACCCTGAAATTTATCTGTACCACCGGCAAACTGCCGG

TTCCGTGGCCGACCCTGGTTACCACCCTGAGCTGGGGTGTTCAGTGTTTTGCACGTTATCC

GGATCACATGAAACAGCACGATTTTTTCAAAAGCGCAATGCCGGAAGGTTATGTTCAAGA

ACGTACCATCTTCTTCAAAGATGACGGCAACTATAAAACCCGTGCCGAAGTTAAATTTGAA

GGTGATACCCTGGTGAATCGCATTGAACTGAAAGGCATCGATTTTAAAGAGGATGGTAATAT

CCTGGGCCACAAACTGGAATATAATGCCATTCATGGCAACGTGTATATCACCGCAGATAAAC

AGAAAAACGGCATCAAAGCAAATTTTGGCCTGAACTGCAATATTGAAGATGGTTCAGTTCA

GCTGGCAGATCATTATCAGCAGAATACCCCGATTGGTGATGGTCCGGTTCTGCTGCCGGATA

ATCATTATCTGAGCACCCAGAGCAAACTGAGCAAAGATCCGAATGAAAAACGTGATCACAT

GGTGCTGCTGGAATTTGTTACCGCAGCAGGTATTACCCTGGGTATGGATGAACTGTATAAAG

GTAGCGGTGGTAGTGGTGGTTCAGGTGGCTCTGGTGGCAGCGGTGGATCCGGTGCAGAAG

CAGCAGCAAAAGAAGCCGCTGCCAAAGAAGCGGCAGCGAAAGAGGCTGCCGCAAAAGA

GGCAGCAGCGAAAGAAGCAGCGGCTAAAGCAGGTTCAGGCGGTTCTGGGGGTTCTGGCG

GTAGCGGAGGCAGTGGCGGTAGTGGCGCTGAGGCAGCCGCTAAAGAAGCTGCGGCAAAA

GAAGCAGCGGCAAAAGAGGCTGCAGCTAAAGAAGCCGCAGCAAAAGAGGCAGCAGCAA

AAGCAGGTAGCGGTGGTTCAGGCGGTAGCGGTGGCTCAGGTGGCAGTGGCGGTTCCATGG

TGTCAAAAGGAGGAACTGTTTACAGGCGTGGTGCCGATCCTGGTAGAGCTGGACGGGGAT

GTGAATGGCCATAAATTCAGCGTTTCAGGTGAAGGTGAGGGCGACGCCACGTACGGAAAA

CTGACACTGAAATTCATTTGCACAACAGGTAAACTGCCTGTGCCTTGCTACACTGGTGACC

ACCTTTGGTTATGGTCTGATGTGCTTTGCTCGCTATCCTGACCACATGAAACAACATGATTT

CTTTAAATCTGCCATGCCTGAAGGCTACGTGCAAGAGCGCACCATTTTTTTCAAAGACGAT

GGGAATTACAAAACACGTGCGGAGGGAAATTCGAGGGCGATACACTGGTTAACCGTATCG

AGCTGAAAGGTATCGACTTCAAAGAGGACGGAAACATTCTGGGTCATAAACTGGAATACA

ACTACAACAGCCATAACGTGTACATCATGGCCGACAAACAAAAAAACGGGATTAAAGTGA

ACTTCAAAATCCGCCACAACATCGAAGATGGCAGCGTGCAGCTGGCCGACCACTATCAAC

AAAACACACCGATCGGCGACGGT

CCTGTACTGCTGCCTGACAACCACTATCTGTCATATCAGAGCGCACTGTCAAAAGATCCTA

ACGAGAAACGCGACCACATGGTTCTGCTGGAATTCGTGACAGCCGCTGGCATTACACTGG

GCATGGACGAGCTGTACAAATAA

#### mVenus-mCherry

ATGAAAGGTGAAGAGCTCTTCACTGGTGTTGTTCCAATTTTGGTTGAATTGGATGGTGATGTTAACGGCCATAAGTTTTCTGTTTCTGGTGAAGGTGAGGGTGATGCTACTTATGGTAAATTGACTTTGAAGTTGATCTGCACCACAGGTAAATTGCCAGTTcccTGGCCAACTTTGGTTACTACTTTAGGTTACGGCTTGCAATGTTTTGCTAGATACCCAGATCATATGAAGCAACACGATTTCTTCAAATCCGCTATGCCAGAAGGTTACGTTCAAGAAAGAACTATCTTCTTCAAGGACGACGGTAACTACAAAACTAGAGCTGAAGTTAAGTTCGAAGGTGATACCTTGGTTAACAGGATTGAATTGAAGGGCATCGATTTCAAAGAGGATGGTAACATTTTGGGTCACAAGTTGGAGTACAACTACAACTCTCATAACGTTTACATTACCGCCGACAAGCAAAAGAATGGTATTAAGGCTAACTTCAAGATCAGGCACAACATTGAAGATGGTGGTGTTCAATTGGCTGATCACTATCAACAAAACACCCCAATTGGTGATGGTCCAGTTTTGTTGCCAGATAACCATTACTTGTCCTACCAGTCCAAGTTGTCTAAAGACCCAAACGAAAAAAGGGATCACATGGTTTTGTTGGAATTTGTTACAGCTGCTTCCGGTGATAACATGGCCATTATCAAAGAATTTATGAGGTTCAAGGTCCACATGGAAGGTTCTGTTAATGGTCACGAATTTGAGATTGAAGGTGAAGGCGAAGGTAGACCATATGAAGGTACTCAAACTGCTAAACTGAAGGTTACAAAAGGTGGTCCATTGCCATTTGCTTGGGATATTTTGTCTCCACAATTCATGTACGGTTCTAAGGCTTATGTAAAACACCCAGCTGATATCCCAGATTACTTGAAGTTGTCATTTCCAGAGGGTTTCAAGTGGGAAAGAGTTATGAATTTCGAGGATGGTGGCGTTGTTACTGTTACTCAAGATTCTTCATTACAGGACGGTGAGTTTATCTACAAGGTTAAGTTGAGAGGTACGAACTTTCCATCTGATGGTCCTGTTATGCAAAAAAAGACTATGGGTTGGGAAGCCTCTTCTGAAAGAATGTATCCAGAAGATGGCGCTTTGAAAGGTGAAATCAAACAAAGGTTGAAATTGAAGGACGGTGGTCATTATGATGCCGAAGTTAAGACTACTTACAAGGCTAAAAAGCCAGTTCAATTACCAGGTGCTTACAACGTCAACATCAAGTTGGATATCACTTCCCACAACGAAGATTACACCATCGTTGAACAATACGAAAGAGCTGAGGGTAGACATTCTACTGGTGGTATGTAA

#### KWK probe

ATGCACCATCACCACCACCACAAGTGGAAGAAGTGGAAAAAAGCATTTAACTCCCATAATGTATATATCACGGCTGATAAACAGAAAAACGGGATCAAAGTCAACTTCACGGTACGTCACAACGTAGAAGACGGTTCCGTTCAGTTAGCGGACCATTATCAACAGAATACTCCCATTGGTGATGGCCCCGTTCTGTTACCTGACAACCACTATCTGTCAACTCAGACAGTATTGTCCAAGGACCCGAATGAGAAACGAGACCACATGGTACTTCACGAATACGTCAACGCAGCAGGTATCACT

CACGGGATGGACGAACTGTATAAAGGAGGTTCAGGTGGCACTATGTCCAAAGGGGAAGAGTTATTCACTGGTGTTGTGCCGATCTTGGTTGAGCTTGATGGTGATGTAAACGGTCATAAATTTTCCGTGCGCGGTGAAGGCGAGGGTGACGCGACTAACGGAAAATTGACCTTGAAGTTCATTTGCACCACGGGAAAGTTACCCGTCCCTTGGCCCACATTAGTGACGACACTTACGTACGGGGTCCAGTGTTTTTCGCGCTACCCGGACCACATGAAGCAGCACGATTTCTTCAAGTCAGCGATGCCCGAGGGCTATGTACAGGAGCGTACTATCTCCTTTAAGGACGACGGCACGTATAAGACTCGTGCAGTTGTAAAATTTGAGGGGGATACCTTGGTCAATCGTATCGAGCTTAAAGGTACCGATTTTAAAGAAGATGGAAATATTCTTGGACACAAACTTGAATACAACAAATGGAAGAAGT

GGAAGAAGGCATAA

